# Mime-seq 2.0: a method to sequence microRNAs from specific mouse cell types

**DOI:** 10.1101/2023.11.06.565867

**Authors:** Ariane Mandlbauer, Qiong Sun, Niko Popitsch, Tanja Schwickert, Miroslava Spanova, Jingkui Wang, Stefan Ameres, Meinrad Busslinger, Luisa Cochella

## Abstract

Many microRNAs (miRNAs) are expressed with high spatiotemporal specificity during organismal development, with some being limited to rare cell types, often embedded in complex tissues. Yet most miRNA profiling efforts remain at the tissue and organ levels. To overcome challenges in accessing the microRNomes from tissue-embedded cells, we had previously developed mime-seq (*<u>mi</u>RNome by <u>me</u>thylation dependent sequencing*), a technique in which cell-specific miRNA methylation in *C. elegans* and *Drosophila* enabled chemo-selective sequencing without the need for cell sorting or biochemical purification. Here, we present mime-seq 2.0 for profiling miRNAs from specific mouse cell types. We engineered a chimeric RNA methyltransferase that is tethered to Argonaute and efficiently methylates miRNAs at their 3’-terminal 2’OH in mouse and human cell lines. We also generated a transgenic mouse for conditional expression of this methyltransferase, which can be used to direct methylation of miRNAs in a cell-type of choice. We validated the use of this mouse by profiling miRNAs from B cells and bone marrow plasma cells.

## Main

MicroRNAs play crucial roles in physiology and development of animals, with individual miRNA expression being highly dependent on the respective cell type. We previously developed mime-seq to retrieve the miRNomes of specific cell types within developing *C. elegans* or *D. melanogaster*^1^. Mime-seq relies on transgenic expression of Hen1 from *Arabidopsis thaliana* (At-Hen1), a methyltransferase involved in the plant miRNA biogenesis pathway^2,3^. With two dsRNA-binding domains, At-Hen1 recognizes products of Dicer cleavage and positions the 3’ 2-nt overhangs of these 21-24 nt long RNA duplexes in the catalytic domain for 2’O-methylation^4^. Transgenic expression of At-Hen1 under cell-specific promoters in worms and flies resulted in efficient cell-specific miRNA methylation. Methylated microRNAs were then selectively detected by performing an oxidation treatment prior to standard small RNA cloning and sequencing. Oxidation by NaIO4 renders the terminal ribose of unmethylated miRNAs non-ligatable while the presence of 2’OMe prevents oxidation and leaves the 3’OH intact and available for adaptor ligation and subsequent sequencing (**Extended Data Fig. 1a**). Because mime-seq proved to be useful for miRNA profiling in *C. elegans* and *Drosophila*, we set out to adapt it for use in mammalian systems, specifically in mice.

To this end, we transfected codon-optimized and Myc-tagged At-Hen1 into human HEK293T cells and assessed miRNA methylation by oxidation of the terminal ribose followed by β-elimination. This treatment removes the 3’- terminal nucleoside of unmethylated but not of methylated miRNAs and causes accelerated migration in high-resolution PAGE that can be visualized by northern blotting (Alefelder, 1998)^5^. In contrast to what we had seen in worms and flies, we were unable to detect 2’O-methylation of endogenous miRNAs, despite detectable expression of At-Hen1 (**Extended Data Fig. 1 b, c**). This was not because the enzyme made in HEK293T cells was inactive, as immunopurified Myc-At-Hen1 from HEK293T was fully active in an *in vitro* methylation assay (**Extended Data Fig. 1d**). This suggested that the difference in At-Hen1 activity in worms and flies versus human cells may reflect different availability of the miRNA duplex generated by Dicer, as in mammals Dicer cleavage is coupled to Argonaute loading^6,7^. To circumvent this, we engineered an alternative methyltransferase to meet the mechanistic requirements of the miRNA biogenesis pathway in mammals.

To develop a suitable methyltransferase, we decided to modify *Mus musculus* HENMT. mHENMT is a methyltransferase specifically expressed in mouse testis^8^, where it is required for piRNA 2’O-methylation at the 3’ terminal ribose^9,10^. Specificity for piRNA methylation is determined by the C-terminal domain of mHENMT that directly interacts with Piwi proteins in which piRNAs are loaded^11^. Contrary to At-Hen1, substrates of mHENMT are single-stranded RNAs of varying lengths^12^. To repurpose mHENMT for miRNA methylation, we removed the C-terminal domain and replaced it with a peptide that binds to the Argonaute (AGO) proteins that load miRNAs (**Fig. 1a**). The GW/TNRC6 family of proteins bind AGO to mediate its repressive function^13^. An 84-amino acid peptide derived from TNRC6B, referred to as T6B peptide, has been shown to effectively interact with all four mouse AGOs^14^. We hypothesized that tethering HENMT to AGO via the T6B peptide might force methylation of loaded miRNAs (**Fig. 1b**). We tested the activity of full-length mHENMT (HENMT^FL^), a C-terminal truncation mutant (HENMT^ΔC^) and the same mutant fused to T6B (HENMT^ΔC^-T6B) by stable integration in mouse embryonic fibroblasts (MEF) and transient transfection in HEK293T cells. Using β-elimination and high-resolution northern blots, we observed that only HENMT^ΔC^-T6B, but not a catalytic mutant or any of the non-T6B versions, methylated miRNAs in MEFs and in HEK293T cells (**Fig. 1c, d, Extended Data Fig. 1e, f**). During these experiments, we noticed variable expression levels of the enzymes, with HENMT^ΔC^ and HENMT^ΔC^-T6B having higher expression level than the others (**Fig. 1d, Extended Data Fig. 1e**). To exclude that expression levels alone determine the large differences in activity, we cloned all HENMTs under a doxycycline inducible TRE3 promoter and integrated them using a lentiviral system into the human cell line RKO. After titrating doxycycline levels to achieve similar expression levels, still only HENMT^ΔC^-T6B showed detectable methylation activity (**Extended Data Fig. 1g, h**). We note that codon-optimized At-Hen1 could never be expressed at levels close to mHENMT. In sum, we designed a new chimeric enzyme that efficiently methylates miRNAs in human (HEK293T and RKO) and mouse (MEF) cells in culture.

**Figure 1.**
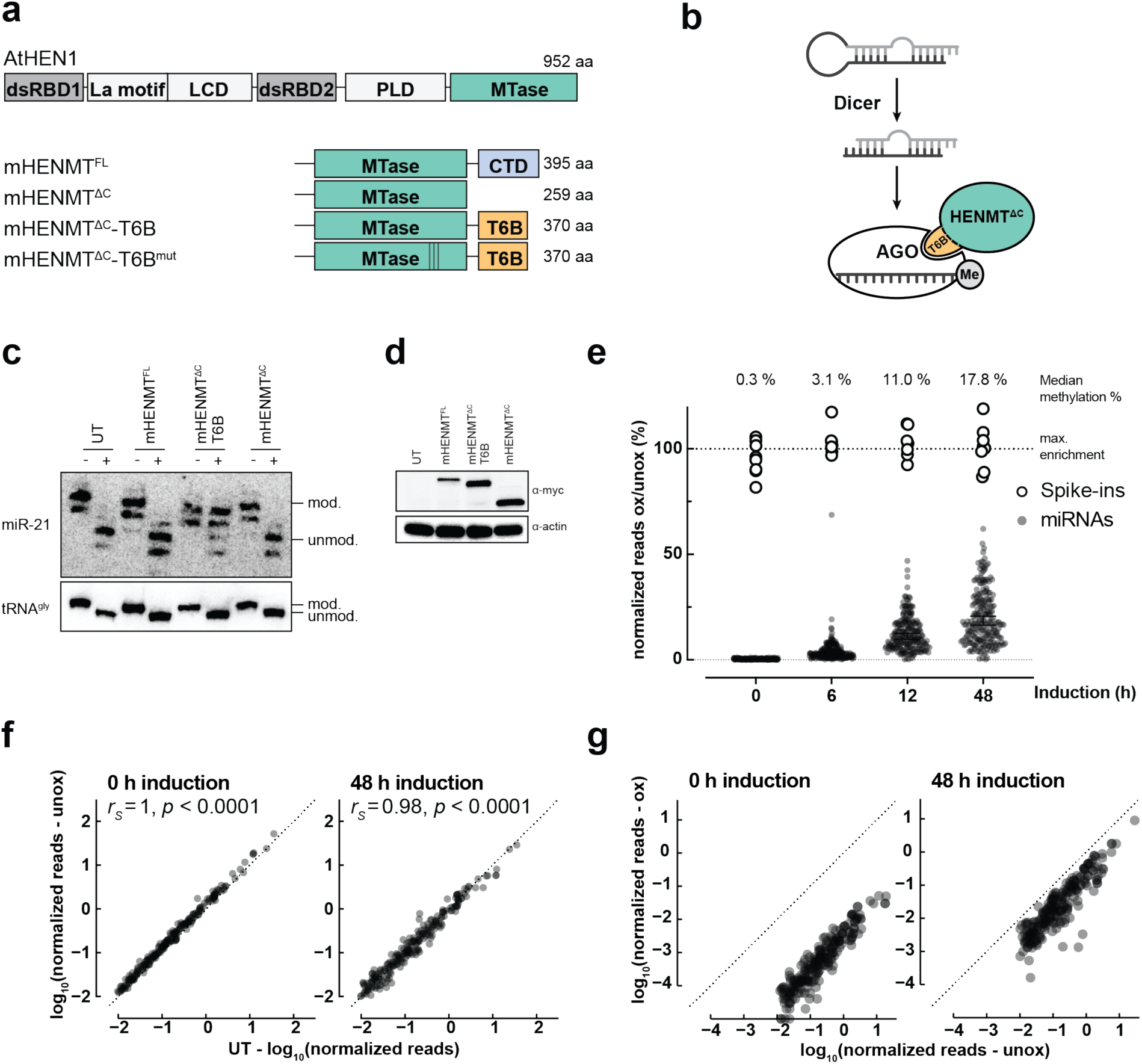
HENMT^ΔC^-T6B, an engineered methyltransferase efficiently methylates mouse and human miRNAs in cultured cells. **(a)** Overview of RNA methyltransferases designed and used in this study. **(b)** Schematic diagram of T6B-dependent tethering of HENMT^ΔC^ to AGO. **(c-d)** Murine embryonic fibroblasts (MEFs) carrying lentivirus-mediated integration of the indicated transgenes under a SFFV promoter. **(c)** Total RNA extracted was subjected to oxidation and β-elimination and the products resolved by high-resolution northern blotting with the indicated probes (tRNA^Gly^) was used to monitor oxidation completion, and as loading control). Position of methylated (mod.) and unmethylated (unmod.) RNAs is indicated. UT: untransduced cells **(d)** Corresponding western blot to detect the various Myc-tagged enzyme variants. Actin served as loading control. **(e-g)** RKO cells with lentivirus-mediated integration of the rtTA3 trans-activator and inducible HENMT^ΔC^-T6B were used for a 48h doxycycline induction time course. Total RNA was extracted at indicated timepoints, methylated spike-ins added, and oxidized and unoxidized samples subjected to small RNA sequencing. MicroRNAs with spike-in-normalized reads <0.01 were removed from the analysis. **(e)** Time-dependent increase of %ox/unox for miRNAs. Fully methylated spike-ins define the maximum possible enrichment. **(f)** Comparison of miRNA levels in untreated samples. Expression of HENMT does not affect miRNA abundance (*rS=*Spearman correlation coefficient, UT: untransduced cells). **(g)** Normalized reads from oxidized vs. unoxidized samples at 0 and 48 h timepoints. The relative abundance of miRNAs is maintained after oxidation (*rS*=Spearman correlation coefficient).

The inducible HENMT^ΔC^-T6B enabled examination of methylation dynamics, by sequencing small RNAs at different times post expression induction (**Fig. 1e, Extended Data Table 1**). We used methylated small RNA spike-ins for normalization^15^, which were added before each sample was split in two, one half was sequenced after the oxidation treatment, and the other half without oxidation (**Extended Data Fig. 2a**). This analysis showed that after 48 hours, 92.6% of miRNAs were methylated ≥5%, which was sufficient for their enrichment after oxidation (**Fig. 1e**). At 48 hours, the miRNAs detected after oxidation represent the top 98.2% of the cumulative miRNA reads in RKO cells, indicating that the physiologically important miRNAs are readily captured by mime-seq 2.0. Importantly, expression of HENMT^ΔC^-T6B did not affect miRNA levels compared to untransduced cells (**Fig. 1f**). Although miRNAs appeared to be methylated with varying rates (**Extended Data Fig. 2b**), perhaps relating to miRNA biogenesis and turnover dynamics^16^, sequencing methylated miRNAs at steady state maintained overall information on relative miRNA expression levels (**Fig. 1g**). Thus, mime-seq 2.0 is semi-quantitative. Altogether, these experiments indicate that HENMT^ΔC^-T6B efficiently methylates miRNAs without substantial bias and should enable the implementation of mime-seq in mammals.

To implement mime-seq 2.0 *in vivo*, we generated the *R26*^LSL-HenT6B/+^ mouse line by CRISPR/Cas9-mediated genome editing in 2-cell embryos^17^ (**Fig. 2a, Extended Data Fig. 3a** and Methods). Upon Cre-mediated deletion of the loxP-Stop-loxP (LSL) cassette, the resulting *R26*^HenT6B/+^ mice give rise to conditional expression of the HENMT^ΔC^-T6B-P2A-eGFP gene from the ubiquitously expressed CAG promoter in the *Rosa26* locus. Hence, specific expression of HENMT^ΔC^-T6B in different cell types can be achieved with appropriate cell type-specific Cre lines. The murine hematopoietic system is ideal for validating the HENMT^ΔC^-T6B-dependent mime-seq *in vivo*. First, because cells are more easily accessible than in other tissues, we can directly compare miRNAs from a purified cell population to a cell mixture, to evaluate the performance of mime-seq. Second, because the different cells of the lineage can be analyzed by flow cytometry, we could quantitatively assess the effect of HENMT^ΔC^-T6B expression in different cell types. This is particularly important given that overexpression of a T6B peptide in the mouse was used as a competitive inhibitor to decrease miRNA function^18^, and that inhibition of miRNA activity causes various defects^19^. We first expressed HENMT^ΔC^-T6B specifically in B cells using the *Cd79a-*Cre line, which induces Cre activity in pro-B and all subsequent stages of B cell development^20^. As shown by flow-cytometric analysis, pro-B, pre-B, recirculating and total B cells in the bone marrow as well as mature B cells in the spleen were minimally affected in *Cd79a*-Cre *R26*^LSL-HenT6B/+^ and *Cd79a*-Cre *R26*^LSL-HenT6B/LSL-HenT6B^ mice compared with control *R26*^LSL-HenT6B/LSL-HenT6B^ mice (**Fig. 2b, c**, **Extended Data Fig. 3b, c**). Moreover, GFP and, by inference, HENMT^ΔC^-T6B were expressed in splenic B cells, but not in T cells of *Cd79a*-Cre *R26*^LSL-HenT6B/+^ and *Cd79a*-Cre *R26*^LSL-HenT6B/LSL-HenT6B^ mice, although the expression was slightly variegated (**Extended Data Fig. 3c**).The presence of the different B cell types in *Cd79a*-Cre *R26*^LSL-HenT6B/LSL-HenT6B^ mice is contrasted by the strong reduction of pre-B cells and absence of all subsequent B cell stages upon conditional Dicer deletion, leading to the loss of all miRNAs in *Cd79a*-Cre *Dicer1*^fl/fl^ mice^21^. We conclude that the expression of HENMT^ΔC^-T6B minimally affects B cell development and is thus compatible with normal miRNA function.

**Figure 2.**
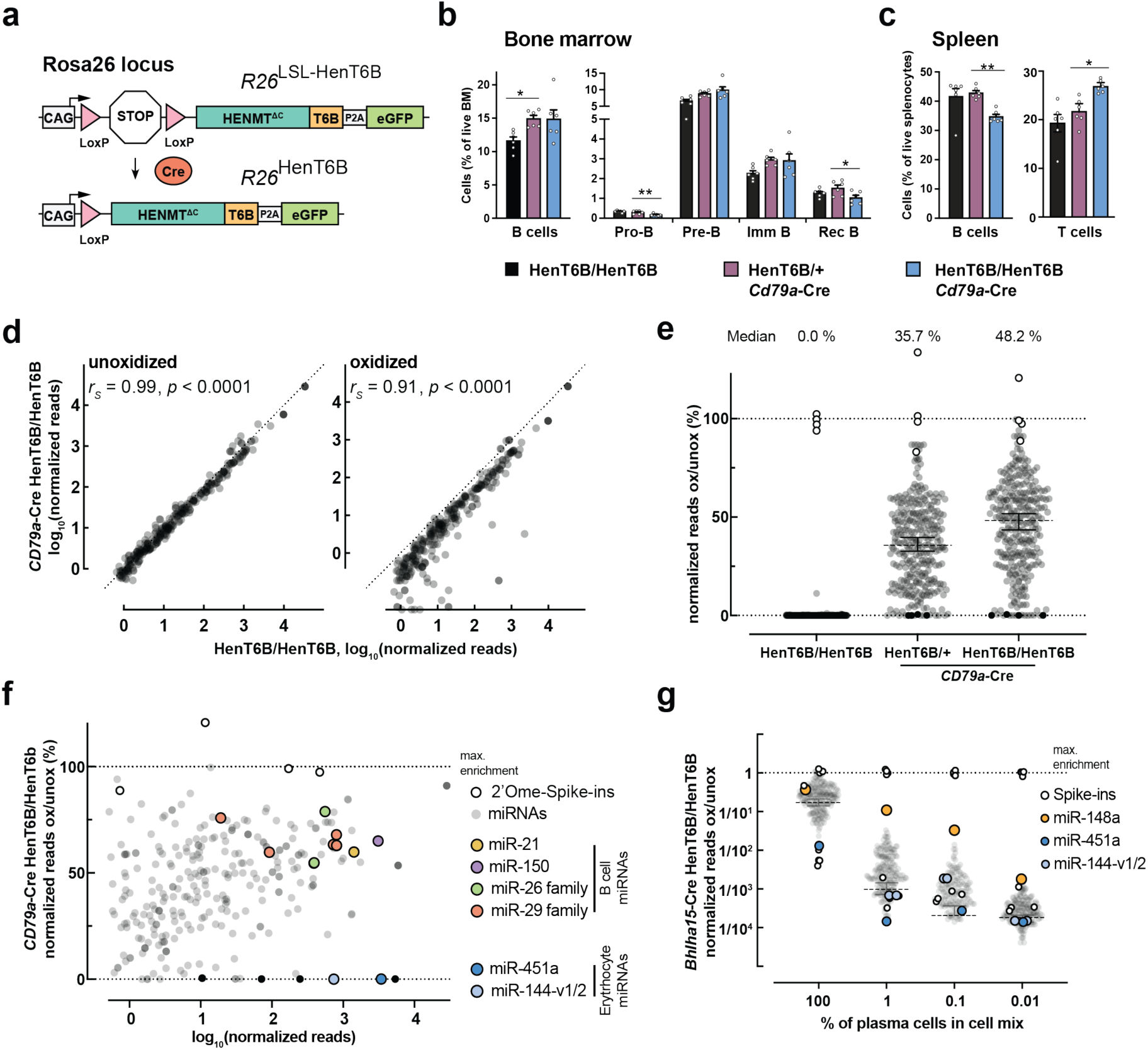
mime-seq using HENMT^ΔC^-T6B methyltransferase allows selective enrichment of cell type specific miRNAs in mice. **(a)** Schematic diagram of the *R26*^LSL-HenT6B^ and *R26*^HenT6B^ alleles. The HENMT^ΔC^-T6B-P2A-eGFP gene is expressed under the control of the CMV enhancer and chicken actin promoter (CAG) from the *R26*^HenT6B^ allele upon Cre-mediated deletion of the loxP-Stop-loxP (LSL) cassette. The CAG promoter in the *Rosa26* (*R26*) locus is known to give rise to ubiquitous expression in different tissues of the mouse. In all further figures, *R26*^LSL-HenT6B/+^ and *R26*^LSL-HenT6B/LSL-HenT6B^ are referred to as HenT6B/+ and HenT6B/ HenT6B, respectively. **(b-c)** Frequencies of different B cell types in the bone marrow **(b)** and spleen **(c)** of 6-8-week-old *Cd79a*-Cre *R26*^LSL-HenT6B/+^ (violet), *Cd79a*-Cre *R26*^LSL-HenT6B/LSL-HenT6B^ (blue) and control *R26*^LSL-HenT6B/LSL-HenT6B^ (black) mice were determined by the flow-cytometric data shown in **Extended Data Fig. 3b,c**. The frequencies of total B, pro-B, pre-B, immature (imm) B, recirculating (rec) B and total T cells are shown as mean values of one independent experiment with SEM. Statistical data (**b**,**c**) were analyzed by one-way ANOVA with Dunnett’s multiple comparison test: \**P* < 0.03, \*\**P* < 0.002. The flow-cytometric definition of the different B cell types is described in Methods. **(d)** Spike-in-normalized small RNA libraries from the same genotypes. Expression of HENMT^ΔC^-T6B does not affect miRNA abundance (unoxidized), and the recovered miRNAs after oxidation largely maintain their relative abundance (oxidized) (*rS*=Spearman correlation coefficient). **(e)** Percentage of recovered reads (ox/unox) are shown. Median level of methylation for every genotype is indicated above. **(f)** Methylation and recovery efficiency as a function of overall abundance in the starting sample. Note enrichment of known B cell miRNAs and depletion of the very abundant miR-451a and miR-144, two miRNAs from contaminating erythrocyte that do not express the methyltransferase. **(g)** Total RNA extracted from *Bhlha15-*Cre *R26*^LSL-HenT6B/LSL-HenT6B^ PCs mixed at indicated ratios with B6 splenocytes. A mix of methylated and unmethylated spike-ins was added and small RNA sequencing libraries prepared from RNA with or without prior oxidation. Reads are normalized to methylated spike ins. Shown are fractions recovered for all sequenced miRNAs with normalized reads >0.5.

To assess the efficiency and specificity of mime-seq *in vivo*, we isolated CD43^−^ mature B cells from the spleen of *Cd79a*-Cre *R26*^LSL-HenT6B/+^, *Cd79a*-Cre *R26*^LSL-HenT6B/LSL-HenT6B^ and control *R26*^LSL-HenT6B/LSL-HenT6B^ mice by immunomagnetic depletion of non-B cells. We extracted total RNA, added a mix of methylated and unmethylated spike-ins (see Methods) and generated small RNA sequencing libraries with and without prior oxidation treatment. After normalization to the methylated spike-ins, we observed that libraries from unoxidized samples showed no significant differences across all genotypes, indicating that expression of HENMT^ΔC^-T6B does not affect miRNA abundance (**Fig. 2d**). Second, we observed that upon oxidation, miRNAs from the mature B cells of control *R26*^LSL-HenT6B/LSL-HenT6B^ mice were depleted to levels comparable with unmethylated spike-ins, indicating that the oxidation treatment efficiently removes non-methylated miRNAs from the pool of sequenced miRNAs (**Extended Data Fig. 4a**). In contrast, HENMT^ΔC^-T6B expressing B cells showed efficient recovery of miRNAs after oxidation, with the homozygous *Cd79a*-Cre *R26*^LSL-HenT6B/LSL-HenT6B^ B cells showing higher levels of methylation/recovery after oxidation than heterozygous *Cd79a*-Cre *R26*^LSL-HenT6B/+^ B cells (**Fig. 2e, Extended Data Fig. 4a**). Importantly, the relative abundance of miRNAs recovered after oxidation was highly similar to the relative abundance of miRNAs in B cells without methylation or oxidation treatment, again showing the semi-quantitative nature of mime-seq (**Fig. 2d**). In addition, miRNAs recovered after oxidation included all miRNA families known to be important for B cell development and homeostasis^22^ (**Fig. 2f**). As the CD43^−^ mature B cells also contained a fraction of contaminating erythrocytes, we could demonstrate that miRNAs known to be absent from B cells, like the erythrocyte-specific miR-451a and miR-144^23^, were fully depleted upon oxidation even if they were present at high levels in the starting sample (**Fig. 2f**). To summarize, mime-seq 2.0 enables specific detection of miRNAs from a cell of interest without strongly influencing miRNA expression and without causing detrimental effects that would be expected from a loss of miRNA function.

As the main advantage of mime-seq is to facilitate miRNA profiling in rare cell populations, it is important to assess its sensitivity. As the *Bhlha15* (Mist1) gene is specifically expressed in plasma cells (PCs) within the hematopoietic system^24^, we used the *Bhlha15*-Cre line for conditional activation of HENMT^ΔC^-T6B expression in splenic PCs. Seven days after immunization with sheep red blood cells, GFP^+^ PCs (CD138^+^TACI^+^) were isolated from the spleen of *Bhlha15*-Cre *R26*^LSL-HenT6B/LSL-HenT6B^ mice by flow-cytometric sorting (**Extended Data Fig. 3d**) and were then mixed at different ratios with wild-type C57BL/6 splenocytes, followed by miRNA sequencing before and after oxidation (**Extended Data Fig. 4b**). The miRNA profiles of PCs (100%) and splenocytes (0.01% PCs) are highly similar with one miRNA exclusively expressed in PCs, miR-148a (**Extended Data Fig. 4c**), which is necessary for PC differentiation and survival^25^. Highlighting the specificity and sensitivity of mime-seq, miR-148a was robustly detected in cell mixes containing 1% and 0.1% PCs, and, although the signal was overall lower, miR-148a was even detected above background in a population with only 0.01% PCs (**Fig. 2g, Extended Data Fig. 4d**). As expected, miRNAs that are expressed in both PCs and splenocytes were enriched to a lower degree than a highly cell-specific miRNA like miR-148a (**Extended Data Fig. 4d, Extended Data Table 2**), but we could still use the full miRNomes to more broadly assess specificity and sensitivity. We first determined a reference PC miRNome composed of 85 miRNAs that account for the top 98% of cumulative reads in the oxidized purified PC population (**Extended Data Fig. 5a, Extended Data Table 2**). We did not use the unoxidized sample because it still contains erythrocytes, given the presence of specific erythrocyte miRNAs that are eliminated after oxidation. We then asked how many of these miRNAs are robustly detected (in the 98^th^ top percentile) after oxidation in the various cell mixes, and also show significant enrichment after oxidation (ox/unox ratio above a cutoff set as the average + standard deviation of the ox/unox ratio of the unmethylated spike-ins, which represent the maximum expected depletion) (**Extended Data Fig. 5b-d**). With these cutoffs, we detected 0 false positives (FPs) in the 1% PC mix and 3 FPs in the 0.1% mix. Moreover, in the 1% PC mix, we recovered 40 of all 85 PC miRNAs, but most importantly we retrieved 5 of the top 5, 9 of the top 10, 22 of the top 25, and 36 of the top 50 PC miRNAs. In the 0.1% mix we still detected 26/50 of the top 50 miRNAs (**Extended Data Fig. 5c-d**). Our results suggest that mime-seq retrieves the most likely physiologically relevant miRNAs (based on abundance and cell-specific expression) in cells that are present in as little as 1/1000 in a mixed cell population.

In summary, mime-seq 2.0 enables sequencing of miRNAs from rare cell types of the mouse. For cell-specific methylation, a conditional *R26*^LSL-HenT6B/LSL-HenT6B^ mouse line was generated that can be crossed with a Cre driver of choice to ultimately induce cell-specific miRNA methylation in the selected cell population. Notably, double heterozygote mice already provide sufficient methylation for the implementation of mime-seq (**Fig. 2, Extended Data Fig. 4a**). Selective cloning and sequencing of 2’O-methylated miRNAs allows semi-quantitative profiling of miRNAs in the cell type of choice. We expect that mime-seq 2.0 will increase the resolution of miRNA profiling efforts to gain the necessary cellular level view to understand miRNA function.

## Acknowledgements

We thank Raphael Manzenreither and Nina Khaldieh from the Ameres group for support with Northern blot experiments and small RNA sequencing libraries. We thank Stephen F. Koniecszny (Purdue University, West Lafayette, USA) for providing the *Bhlha15*^Cre/+^ mouse line, C. Theussl’s team for generating the *R26*^LSL-HenT6B/+^ mouse line, K. Aumayr’s team (IMP, Vienna) for flow-cytometric sorting, the VBCF NGS facility for all Illumina sequencing. This research was supported by Boehringer Ingelheim, and NSF CAREER Award 2238425 to L.C. Research in the Ameres group is supported by the European Research Council (ERC-CoG-866166) and the Austrian Science Fund FWF (SFB-F8002).

## Author contributions

A.M. designed the enzyme variant, performed experiments and analyzed the data, unless otherwise noted; Q.S. characterized the *R26*^LSL-HenT6B/+^ mouse by performing flow-cytometric analyses; N.P. and J.W. performed all bioinformatic analyses; T.S. generated the *Bhlha15*-Cre *R26*^LSL-HenT6B/LSL-HenT6B^ mice and isolated the plasma cells for the mixing experiment; M.S. performed the in vitro methylation assay; M.B designed the strategy for generating the *R26*^LSL-HenT6B^ allele and planned the mouse work; A. M., S.A. and L.C. planned the project, designed the experiments and wrote the manuscript together with M.B.

## Extended Data Figures

**Extended Data Figure 1.**
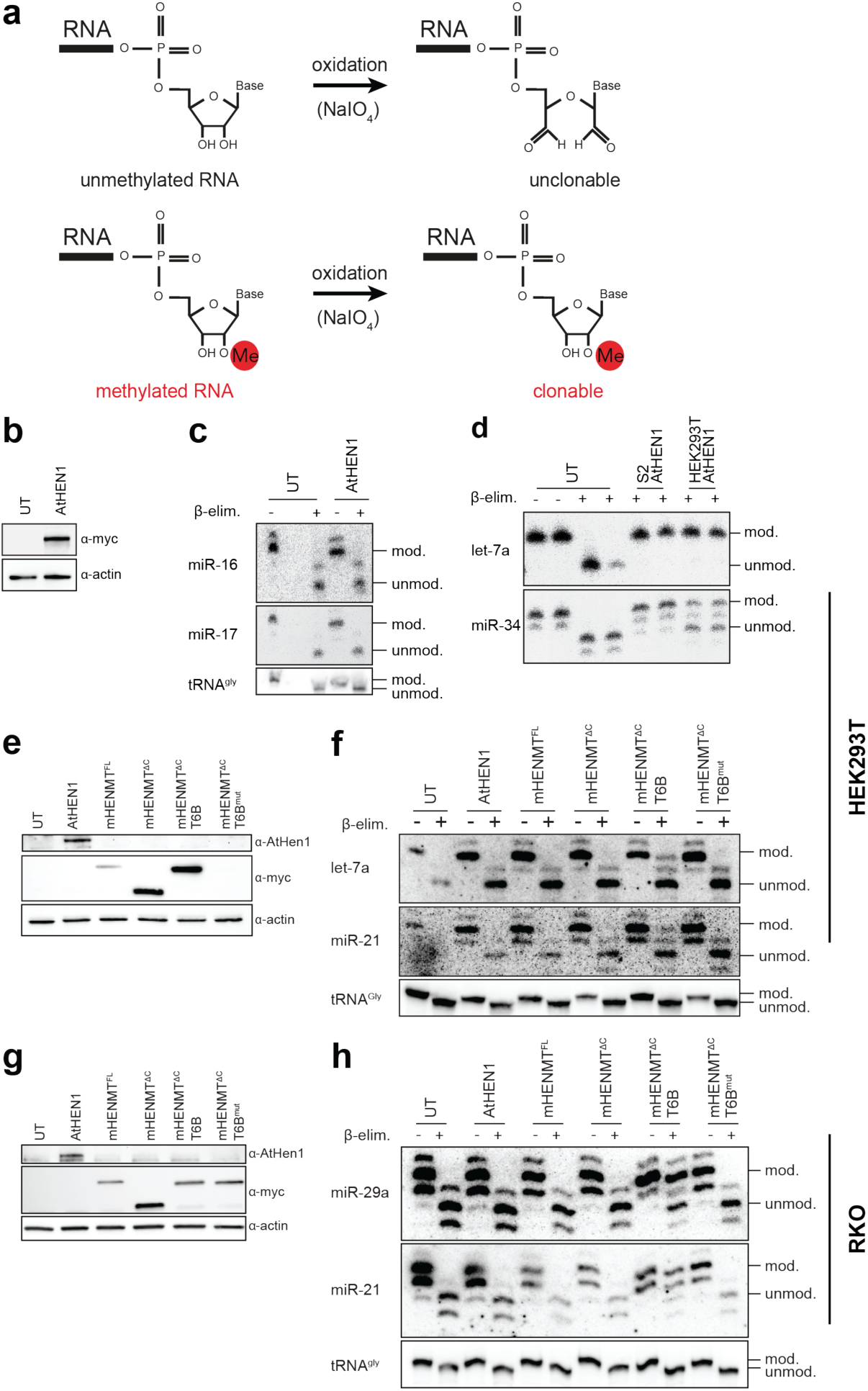
HENMT^ΔC^-T6B but not AtHen1 efficiently methylates mouse and human miRNAs in cultured cells. **(a)** Schematic of the reaction enabling selective adapter ligation of methylated RNA. Unmethylated small RNAs are oxidized by NaIO4, 2’Omethylated miRNAs are protected and retain a free 3’OH, required for 3’ adapter ligation. **(b-f)** HEK293T cells transfected with a construct for expression of codon optimized and Myc-tagged At-Hen1. **(b)** Western blot for Myc-tagged At-Hen1 from the HEK293T cells. Actin served as loading control. **(c)** Total RNA from the HEK293T cells was subjected to oxidation and β-elimination and the products resolved by high-resolution northern blotting with the indicated probes (tRNA^Gly^ was used to monitor oxidation completion, and as loading control). **(d)** *In vitro* methylation assay. FLAG-tagged At-Hen1 was immunopurified upon expression in S2 cells, or Myc-tagged At-Hen1 from HEK293T cells and incubated at 37°C with radiolabeled dme-let-7 or dme-miR-34 duplex RNAs with 2-nt 3’ overhangs. Incubation without enzyme served as negative control. Methylation was assessed by β-elimination and high-resolution PAGE. **(e)** Western blots to monitor expression of the various enzymes in HEK293T cells transfected with indicated constructs. **(f)** Total RNA extracted from HEK293T cells transiently transfected with indicated constructs, and untransfected control (UT) was treated as in (Fig. 1c). **(g-h)** RKO cells transduced with indicated constructs under a doxycycline inducible TRE3 in cells that have integrated the rtTA3 transactivator. **(g)** Western blots to monitor expression of the various enzymes with titrated doxycycline concentrations. **(h)** Northern blots as described in (c).

**Extended Data Figure 2.**
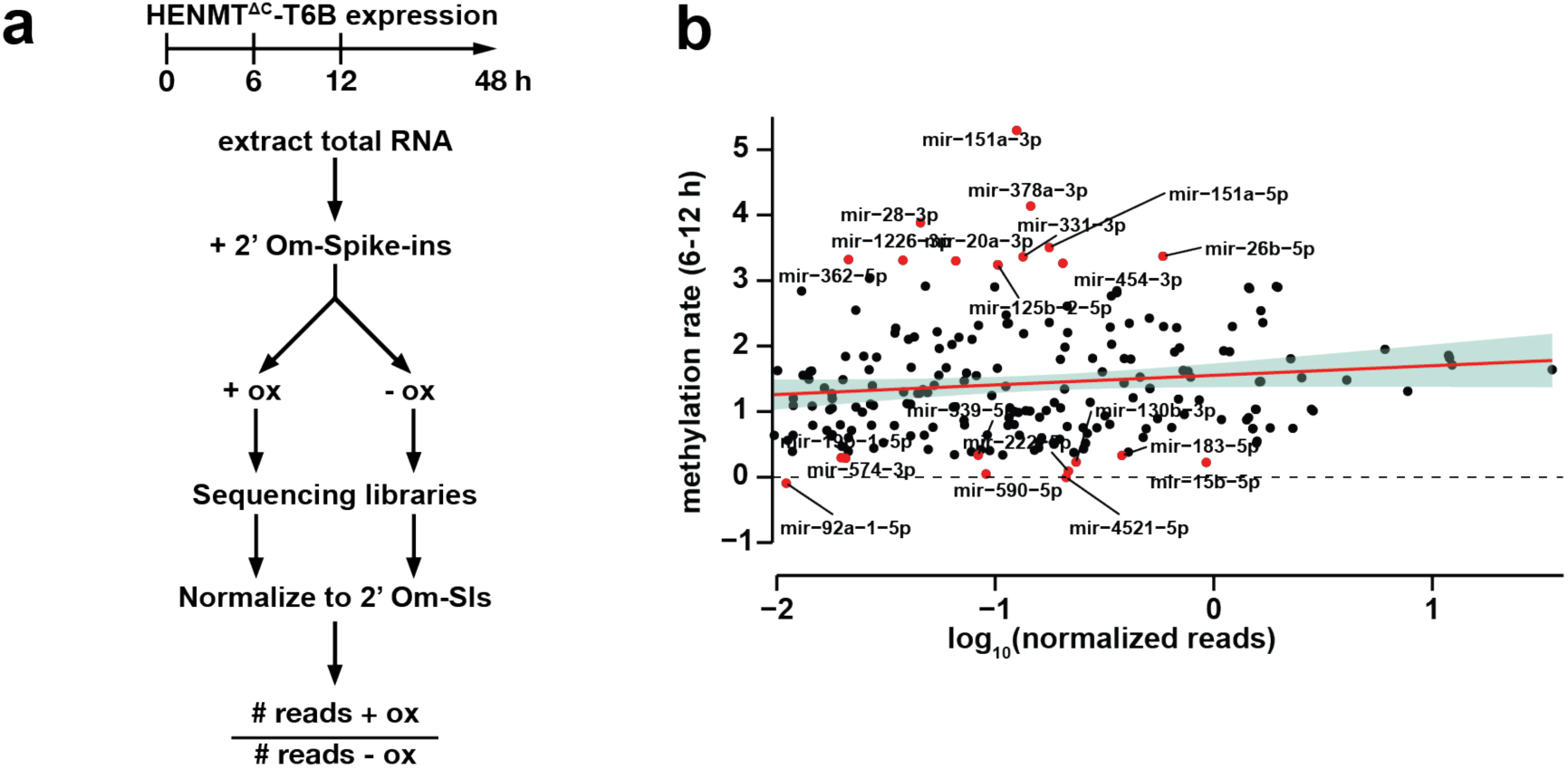
Analysis of methylation kinetics in HENMT^ΔC^-T6B expressing RKO cell line. **(a**) Schematic overview of the timecourse experiment. Total RNA was extracted from cells after 0-48 h of HENMT^ΔC^-T6B expression. After addition of methylated and unmethylated spike-ins samples are treated with or without NaIO4 followed by generation of sequencing libraries. To investigate what fraction of miRNAs are methylated, we calculated the ratio of normalized reads from oxidized to unoxidized samples. **(b)** Methylation rate, calculated as the slope of % ox/unox between 6 and 12h of induction, does not correlate with miRNA abundance. miRNAs with the top and bottom 5th percentile methylation rates are colored.

**Extended Data Figure 3.**
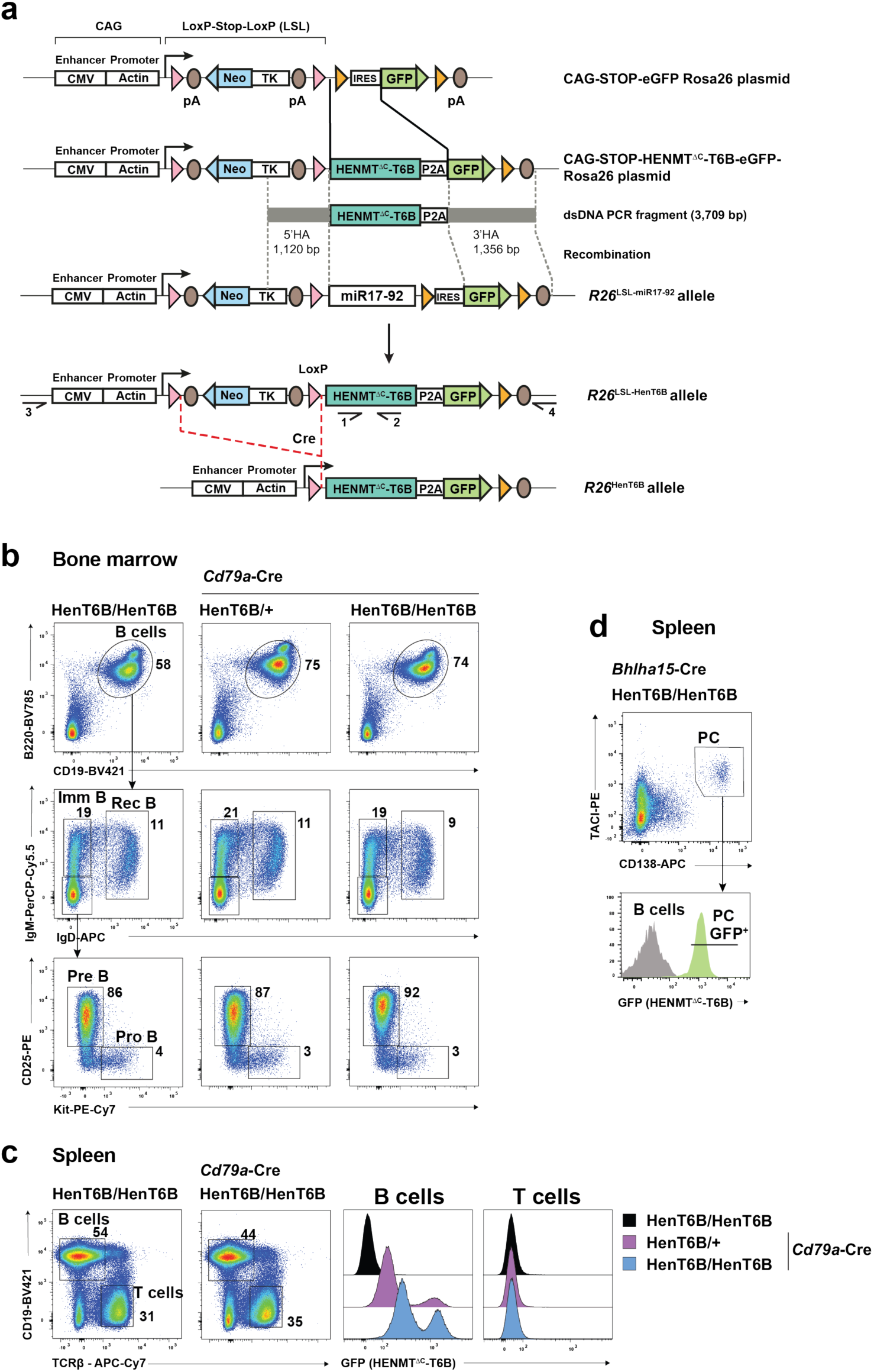
Generation of the *R26*^LSL-HenT6B^ allele and analysis of B cell development in *Cd79a*-Cre *R26*^LSL-HenT6B/ LSL-HenT6B^ mice. **(a)** Generation of the *R26*^LSL-HenT6B^ allele. The HENMT^ΔC^-T6B cDNA was cloned in the CAG-STOP-eGFP-Rosa26 TV plasmid by replacing the IRES sequence with the HENMT^ΔC^-T6B cDNA fused in frame via a P2A peptide to the eGFP coding sequence to generate the CAG-STOP-HENMT^ΔC^-T6B-eGFP-Rosa26 plasmid. A 3,709-bp PCR fragment containing a 1,120-bp 5’ homology arm (HA) and a 1,356-bp 3’ HA was used as double-stranded DNA repair template together with two sgRNAs to replace the miR17-92-IRES DNA sequences of the *R2*6^LSL-miR17-92^ allele^26^ by CRISPR/Cas9-mediated genome editing in mouse 2-cell embryos, resulting in the *R26*^LSL-HenT6B^ allele. Cre-mediated deletion of the loxP-Stop-loxP (LSL) cassette leads to the expression of the HENMT^ΔC^-T6B-P2A-eGFP gene from the ubiquitously transcribed CAG promoter of the *R26*^HenT6B^ allele. *LoxP* and *frt* sites are indicated by red and yellow arrowheads, respectively. The herpes simplex virus thymidine kinase (TK) promoter drives expression of the neomycin (Neo) resistance gene. Arrows indicate primers 1 and 2 used for genotyping of the *R26*^LSL-HenT6B^ allele and primers 3 and 4 used for genotyping of the wild-type *R26* allele (see Methods). pA, polyadenylation sequence. **(b)** Flow-cytometric analyses of the indicated B cell types in the bone marrow of *Cd79a*-Cre *R26*^LSL-HenT6B/+^, *Cd79a*-Cre *R26*^LSL-HenT6B/LSL-HenT6B^ and control *R26*^LSL-HenT6B/LSL-HenT6B^ mice at the age of 6-8 weeks. The percentage of cells in the indicated gates is shown. One of 3 independent experiments is shown. **(c)** Flow-cytometric analysis of B and T cells in the spleen of *Cd79a*-Cre *R26*^LSL-HenT6B/+^, *Cd79a*-Cre *R26*^LSL-HenT6B/LSL-HenT6B^ and control *R26*^LSL-HenT6B/LSL-HenT6B^ mice at the age of 6-8 weeks. The percentage of B and T cells is shown next to the respective gate (left). The GFP expression of B and T cell of the indicated genotypes is shown as a histogram (right). One of 3 independent experiments is shown. **(d)** Flow-cytometric sorting of plasma cells from the spleen of *Bhlha15*-Cre *R26*^LSL-HenT6B/LSL-HenT6B^ mice at day 7 after immunization with sheep red blood cells. The sorting gates used for the isolation of plasma cells (CD138^+^TACI^+^) are indicated.

**Extended Data Figure 4.**
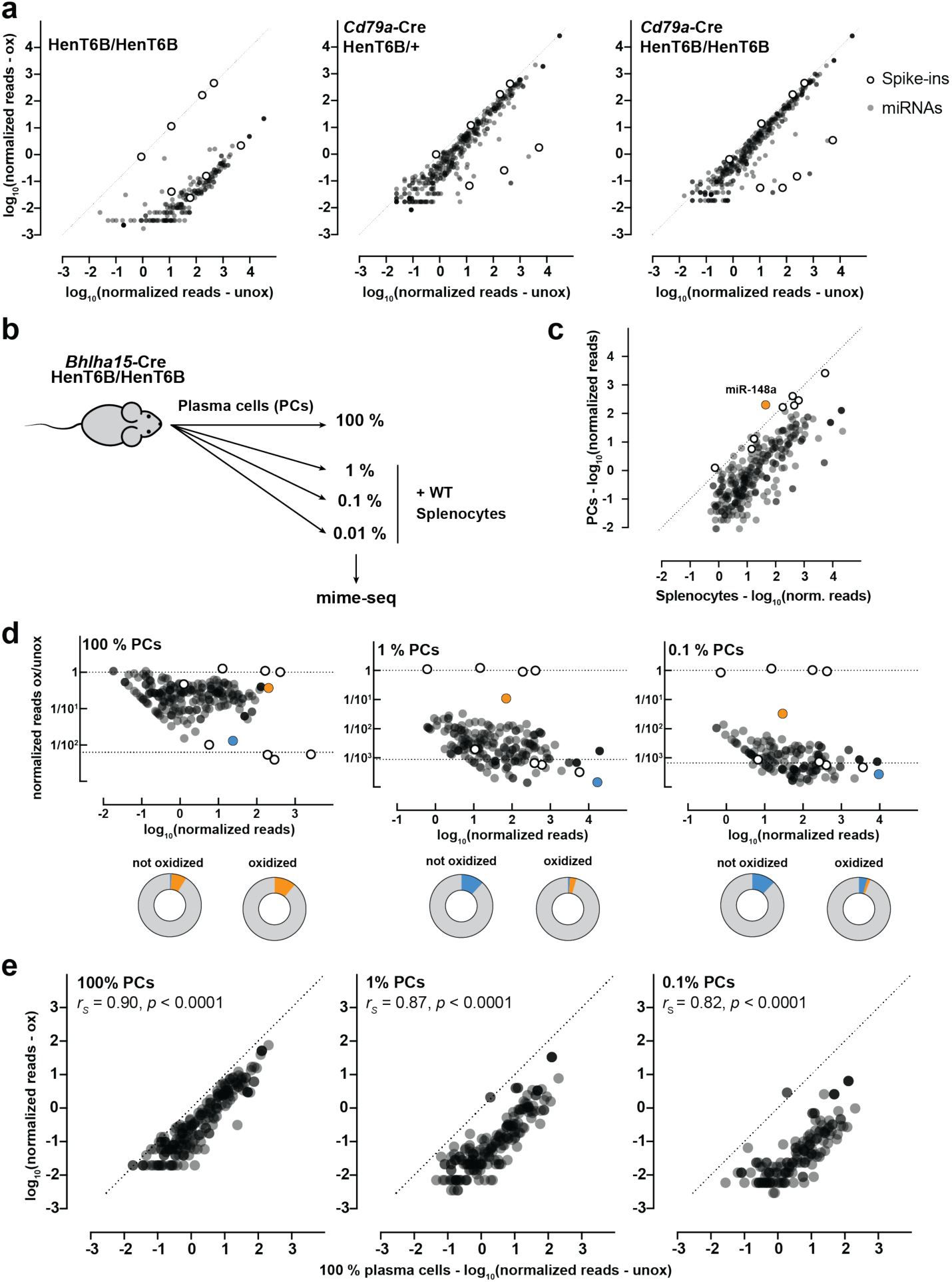
Mime-seq identifies cell-specific miRNAs with high sensitivity while maintaining relative abundance of miRNAs. **(a)** Total RNA was extracted from B cells of the indicated genotypes (see Methods for purification strategy), mixed with four methylated and four unmethylated RNA spike-ins and subjected to mime-seq. Libraries were normalized to methylated spike-ins. Normalized reads of oxidized vs. unoxided libraries are plotted for every miRNA with > 0.5 norm. reads. Note the depletion of unmethylated miRNAs (similar to the four unmethylated spike-ins) and the strong enrichment when HenT6B is expressed (close to the four methylated spike-ins)**. (b)** Schematic overview of the mixing experiment to test the sensitivity of mime-seq. Plasma cells (PCs) were isolated from the spleen of immunized *Bhlha15-*Cre *R26^HenT6B/HenT6B^*mice by flow-cytometric sorting (**Extended Data** Fig. 3d) and were then mixed at the indicated ratios with total splenocytes of wild-type (WT) C57BL/6 mice. Mime-seq was performed with the different cell mixes. **(c)** Comparison of unoxidized, spike-in normalized small RNA libraries from 100% PCs vs 99.99% splenocytes (0.01% PCs) showed that miR-148a is exclusively expressed in PCs. **(d)** Fractions recovered for all sequenced miRNAs with normalized reads >0.5 (PC mixing ratios indicated above). Pie charts below show proportions of miRNA reads from oxidized and unoxidized samples, highlighting the enrichment of miR-148 (orange) upon oxidation even from samples with 0.1% PCs, as well as the depletion of miR-451 (blue) from contaminating erythrocytes. **(e)** Oxidized libraries from samples with the indicated mixing ratios vs. normalized reads from unoxidized pure PCs. The relative abundance of miRNAs present in PCs is maintained after oxidation, showing that mime-seq remains semi-quantitative down to 0.1% PC (*rS*=Spearman correlation coefficient).

**Extended Data Figure 5.**
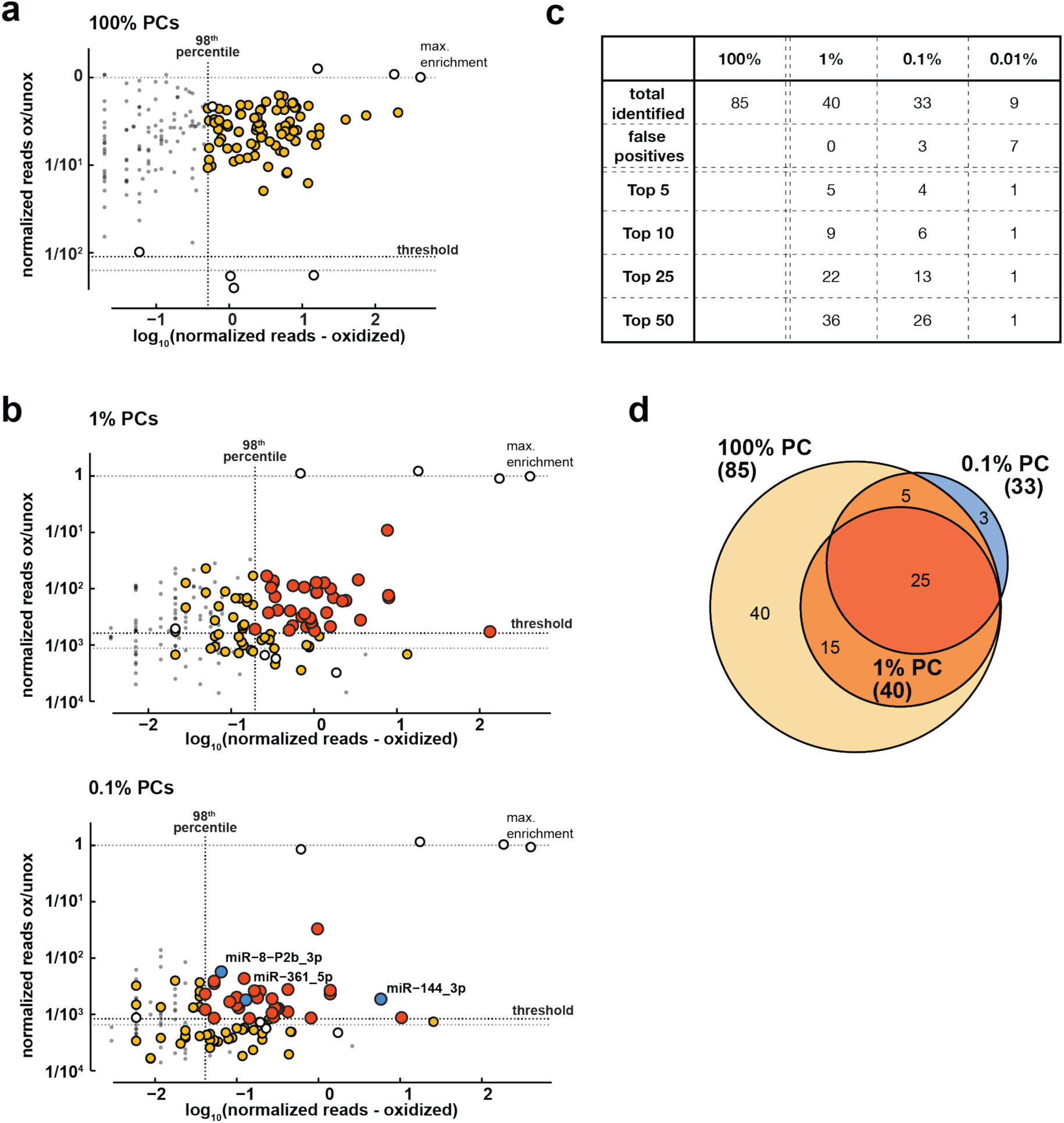
HENMT^ΔC^-T6B mime-seq recovers cell-specific miRNAs *in vivo*. **(a-b)** Identification of miRNAs confidently expressed in PCs. Black dotted lines indicate the thresholds use to define miRNAs expressed in PCs. We used an abundance cutoff (vertical line) for miRNAs that are in the top 98^th^ percentile of cumulative reads, and an enrichment after oxidation cutoff (horizontal line) calculated as the average + standard deviation of the ox/unox ratio of the four unmethylated spike-ins. Shown are fractions recovered for all sequenced miRNAs with normalized reads >0.5 (PC mixing ratios indicated above). Reads originating from isomiRs are summed. **(a)** miRNAs identified in 100% PC (yellow). **(b)** miRNAs identified in 1% or 0.1% PC (orange). False positives (blue) are defined as miRNAs that pass the set thresholds but are not identified in the 98^th^ top percentile of the 100% PC sample. **(c)** Summary table of the most highly abundant PC miRNAs recovered in 1%, 0.1% or 0.01% PCs. **(d)** Venn-diagram showing overlap between different samples. Colors as described above.

## Materials and Methods

### Vectors

All HENMT and At-Hen1 constructs were cloned into a modified lentiviral pLX303 vector under a SFFV promoter. Sequences for HENMT-T6B and codon-optimized At-Hen1 were ordered as gene blocks, modified constructs shown in Fig. 1a were generated using Gibson assembly or side-directed mutagenesis. To generate inducible expression vectors, SFFV promoter was replaced with a TRE3 promoter by Gibson assembly. All constructs were also cloned into a minimal pCS2 vector under a CMV IE94 promoter for transfection. All plasmids used in this study are provided in **Extended Data Table 3**. Lentiviral HENMT^ΔC^-T6B plasmid will be made available through Addgene.

### Cell maintenance

Cell lines were maintained in DMEM (HEK293T) or RPMI (RKO) supplemented with 10% FBS (Avantor, 89510186), 4mM l-glutamine (Gibco, 25030081), 1mM Sodium Pyruvate (Sigma, S8636) and 25 mM HEPES (Sigma, H4034-500G) at 37°C, 5% CO2.

### Transfection

6x10^5^ cells per well were seeded the day before transfection in a 6-well plate. 1 µg plasmid DNA was transfected with polyethylenimine (PEI, Polysciences, 23966) at a ratio of 1:3 (µg DNA: µg PEI) in 200 µl Opti-MEM (Gibco, 31985070). Media was replaced after 24h. After 48h cells were washed with ice-cold PBS and harvested either in RIPA buffer (150 mM NaCl, 1 % IGEPAL CA-630, 0.5 % Sodium deoxycholate, 0.1 % SDS, 50 mM Tris pH 8) for Western blot experiments or in TRIzol (Invitrogen, 15596026) for RNA extraction. Samples were stored at -80°C.

### Lentivirus production and target cell transductions

Commercial LentiX (Takara, 632180) at 70% confluency were transfected with a mix of Eco envelope plasmid (Cell Biolabs, 320026), pCMVR8.74 (Addgene, 22036) and transfer plasmid of interest. Supernatant, containing VLPs, was harvested after 48h, 56h and 72h, filtered (0.45 µm) and used for target cell transduction in the presence of 4 ug/ml polybrene (Sigma). Blasticidin (InvivoGen, ant-bl-05) was used at a concentration of 5 µg/ml for selection of integrated constructs.

### Mouse related methods

The following mice were maintained on the C57BL/6 genetic background: Cd79aCre/+ mice^1^, homozygous miR-17-92 transgenic mice^2^ (here referred to as R26^LSL-miR17-92/LSL-miR17-92^ mice), and Bhlha15Cre/+ mice (generated in the lab of Stephen F. Koniecszny, Purdue University, West Lafayette, USA). Cd79aCre/+ and Bhlha15Cre/+ mice are referred here as *Cd79a*-Cre and *Bhlha15*-Cre mice, respectively. All animal experiments were carried out according to valid project licenses, which were approved and regularly controlled by the Austrian Veterinary Authorities.

### Generation of the ^R26LSL-HenT6B^ allele

For generating the *R26*^LSL-HenT6B^ allele, the HENMT^ΔC^-T6B cDNA was first cloned in the CAG-STOP-eGFP-Rosa26 TV plasmid (Addgene, 15912) by replacing the IRES sequence with the HENMT^ΔC^-T6B cDNA fused in frame via a P2A peptide to the eGFP coding sequence to generate the CAG-STOP-HENMT^ΔC^-T6B-eGFP-Rosa26 plasmid (**Extended Data Fig. 3a**). A 3,709-bp long DNA fragment was PCR-amplified from the CAG-STOP-HENMT^ΔC^-T6B-eGFP-Rosa26 plasmid with an upstream primer (5’-CTGGCACTTCTTGGTTTTCC-3’) and downstream primer (5’-GCTGCATAAAACCCCAGATG-3’) (**Extended Data Fig. 3a**). The *R26*^LSL-HenT6B^ allele was generated by CRISPR/Cas9-mediated genome editing in mouse 2-cell embryos (2C-HR-CRISPR)^3^. For this, 2-cell embryos of the *R2*6^LSL-miR17-92/LSL-miR17-19^ genotype (C57BL/6) were injected with Cas9 protein, two appropriate sgRNAs (linked to the scaffold tracrRNA) and the double-stranded 3,709-bp DNA repair template to generate the *R26*^LSL-HenT6B^ allele (**Extended Data Fig. 3a**). Correct targeting of the *R26*^LSL-HenT6B^ allele was verified by DNA sequencing of respective PCR fragments. The *R26*^LSL-HenT6B^ allele was genotyped by amplification of a 334-bp PCR fragment from the HENMT^ΔC^ insert with primer 1 (5’-ATGCCAAGCTCCTAAAGCTG-3’) and primer 2 (5’- GGGTTGAATTCAGCATTTGG-3’). In a separate PCR, the wild-type *R26*^+^ allele was genotyped by amplification of a 170-bp PCR fragment with primer 3 (5‘-CTCTTCCCTCGTGATCTGCAACTCC-3’) and primer 4 (5’-TCCCGACAAAACCGAAAAT-3’).

### Definition of the cell types by flow cytometry

The different hematopoietic cell types of the mouse were identified by flow cytometry as follows: pro-B (B220^+^CD19^+^Kit^+^CD25^−^IgM^−^IgD^−^), pre-B (B220^+^CD19^+^Kit^−^CD25^+^IgM^−^IgD^−^), immature B (B220^+^CD19^+^IgM^hi^IgD^−^), recirculating B (B220^+^CD19^+^IgM^+^IgD^hi^), plasma cells (CD138^+^TACI^+^), total B cells (B220^+^CD19^+^) and total T cells (CD3^+^TCRΔ^+^). Flow-cytometric analysis was performed on the LSRFortessa (BD Biosciences) machine and flow-cytometric sorting of plasma cells on the FACSAria III (BD Biosciences) machine. FlowJo Software (Treestar) was used for data analysis. CD43^−^ B cells were enriched from the spleen by immunomagnetic depletion of non-B cells with CD43-MicroBeads (Miltenyi Biotec). The following monoclonal antibodies were used for flow-cytometric analysis of mouse lymphoid organs from 4-12 week-old mice: B220/CD45R (RA3-6B2), CD3 (17A2), CD4 (GK1.5), CD8a (53-67), CD11b/Mac1 (M1/70), CD19 (1D3), CD21/CD35 (7G6), CD23 (B3B4), CD25 (PC61.5), CD117/Kit (ACK2), CD138 (281-2), CD267/TACI (8F10), IgD (11-26C), IgM (II/41) and TCRβ (H57-597) antibodies.

### Immunization

Sheep red blood cells were washed in PBS and resuspended at 10^9^ cells/ml followed by intraperitoneal injection of 100 ml into an adult mouse. Plasma cells from *Bhlha15*-Cre *R26*^LSL-HenT6B/LSL-HenT6B^ mice were isolated by flow-cytometric sorting at day 7 after immunization.

### Western blot

Cells were lysed in RIPA buffer. Protein concentration was determined by BCA (Thermo Scientific, 23227). After addition of 6x loading dye (200 mM Tris pH 6.8, 10% SDS, 60% Glycerol, 0.036% Bromphenol Blue, 5% BME), 15-30 µg of protein were denatured 5 min at 95°C, separated on a 4-20% gradient gel (BioRad, 4561093) and transferred to a nitrocellulose membrane (BioRad). After blocking for 30 min (5% milk in Tris-buffered saline, 0.1% Tween20 - TBST), membrane with primary antibody was incubated overnight at 4°C. The next day membranes were washed three times with TBST and incubated with HRP-conjugated secondary antibodies (5% milk in TBST) for 1 h at room temperature (RT). Membranes were washed three times with TBST and imaged with Clarity Western ECL (BioRad, 1705060). All blots were imaged with a ChemiDoc (BioRad) or Odyssey XF (LI-COR) imager. For detection of Myc-tagged HENMT constructs and Myc-At-Hen1 mouse monoclonal anti-Myc tag antibody, clone 4A6 (Merck Millipore, 05-724) at 1:2,000-1:5,000 dilution and rabbit anti-AtHen1 antibody (Agrisera, AS15 3095) at 1:1,000 dilution were used. For detection of Actin rabbit anti-Actin antibody (Sigma, A2066) was used at 1:2,000 dilution. Secondary HRP-coupled antibodies, anti-rabbit IgG HRP-linked antibody (Cell Signaling Technology, 7074) and anti-mouse IgG HRP-linked antibody (Cell Signaling Technology, 7076) were used at 1:2,000 dilution.

### RNA extraction and preparation

Samples were lysed in TRIzol. For frozen cell pellets, samples were thawed on ice before addition of 1 ml TRIzol. 200 µl chloroform (Fisher Chemical, C298-500) was added per 1 ml of TRIzol, samples vortexed and incubated for 3 min at RT. Samples were then centrifuged for 15 min at 4°C (12,000 x g) to promote separation of organic (bottom) and aqueous phase (top). Clear phase on top was transferred to a new 1.5 ml tube and 1 µl glycogen (20 mg/ml, Sigma, missing) added. RNA was precipitated by addition of 1 volume isopropanol (Fisher Chemical, A416-1). Samples were vortexed, incubated 5 min at RT and pelleted by centrifugation for 10 min at 4°C (12,000 x g). Pellets were washed with 80% ice-cold EtOH. For resuspension of RNA, pellets were centrifuged and air dried for 5 min before addition of RNase-free H2O (Ambion). RNA was quantified using the Qubit BR kit (Invitrogen, Q10210). For northern hybridization experiments, 5-15 µg RNA was subjected to oxidation followed by β-elimination per lane. For small RNA library preparation, when possible >3 µg RNA was mixed with methylated and unmethylated spike-ins^4^ (sequences are provided in supplementary table X) before sample was split in half for oxidation with and without NaIO4. Where indicated, libraries were generated from the maximum amount of RNA obtained, e.g. from sorted plasma cells we only retrieved 340 ng of RNA and still were able to produce good quality libraries.

### Northern blot

**Oxidation.** For a typical 20 µl reaction, 5-15 µg RNA were used as input with or without 2 µl NaIO4 (50 mM) and 4 µl 5x borate buffer (300 mM boric acid/borax, pH 8.6). Reactions were carried out at RT for 30 min. **β-elimination.** 1 µl NaOH (1 M) was added to a final concentration of 50 mM and incubated at 45°C for 90 min. Reactions were filled up with H2O to 300 µl with 300 mM NaOAc (pH 5.2, Ambion). **Northern Blotting.** Samples were precipitated using 900 µl EtOH using glycogen as carrier and incubated for 1.5h at -20°C before pelleting for 30 min at 4°C with 20,000 x g. After a wash step using 80 % EtOH sample pellets were resuspended in 10 µl of formamide loading dye (Invitrogen, AM8547) and loaded on a 15% urea PAGE (National Diagnostics, EC-833). Northern blots were performed as previously described^5^ with minor modifications. Using bromophenol blue and xylene cyanole as reference, the gel was cut in two. After semidry transfer (BioRad) onto Hybond NX membranes (Cytiva, RPN303T), the lower part was used for hybridization and visualization of miRNAs, the upper part for the tRNA used as loading control and as a control that the oxidation reaction proceeded to completion. RNA was UV crosslinked three times with 120 mJ/cm^2^ each, followed by chemical crosslinking using methylimidazole/EDC^6^. Membranes were prehybridized with Church buffer (1 mM EDTA, 0.5 M Na2HPO4/NaH2PO4, 7 % SDS) for 1 h at 65°C in a hybridization oven. Probes were synthesized as ssDNA oligos and 5’ ^32^P-radiolabeled with T4 polynucleotide-kinase (NEB) and γ-^32^P-ATP (6000 Ci/mmol, Perkin Elmer). All probe sequences used in this study are provided in Extended Data Table 1. Unincorporated nucleotides were removed by G25 column purification (Cytiva, 27532501). 25 pmoles radioactively labeled probes were added. Temperature was lowered to 37°C and membranes incubated overnight. Membranes were washed at 37°C three times with 1x SSC + 0.1% SDS for 10 min before exposure to a storage phosphor screen (Cytiva). All imaging was performed with a Typhoon phosphorimager (Cytiva).

### In vitro methylation assay

*In vitro* methylation assays were performed using immunopurified Myc-AtHEN1 from transiently transfected HEK293T cells or FLAG-Myc-AtHEN1 from stably expressing S2 cells. Immunoprecipitations were performed as previously described^7^ using Protein G Dynabeads (LifeTechnologies) coupled to Myc 4A6 antibody (Merck Millipore, 05-724) or FLAG M2 monoclonal antibody (Sigma, F1804). All RNA substrate sequences can be found in Extended Data Table 1. For miRNA substrate preparation, guide strand was 5’ ^32^P-radiolabeled using T4 polynucleotide-kinase and unincorporated nucleotides removed by G25 column purification. After PAGE purification, labeled guide strands were annealed to 5′ phosphorylated miR* strands. Purified methyltransferases were incubated with labeled miRNA substrates in standard RNAi reactions containing S-adenosylmethionine. Following phenol/chloroform extraction, RNA was subjected to oxidation and β-elimination and run on a 15% denaturing PAGE. Dried gels were exposed to a storage phosphorscreen and imaged using a phosphorimager.

### Small RNA libraries

**Oxidation.** Performed as described above. For small RNA libraries, no β-elimination was performed. After oxidation, samples were purified by EtOH precipitation as described and resuspended in 6 µl H2O. **3’ adapter ligation and purification.** Barcoded 3’ adapters (see Extended Data Table 1) were ligated using K227Q truncated T4 RNA ligase 2 (NEB, M0351L) at a concentration of 0.5 µM adapter and in the presence of 25% PEG8000 containing a homemade 10 x ligation buffer (0.5 M Tris pH 7.8, 0.1 M MgCl2, 0.1 M DTT) overnight at 4°C. As reference for later PAGE purification, independent ligation reactions were also set up for an 18-mer and 30-mer ssRNA oligos. 3’ adapter ligated small RNAs were loaded in formamide loading buffer on a 15% urea PAGE. After visualizing RNA with SYBR Gold (1:7,500; Invitrogen, S11494) in 0.5x TBE, gel fragments spanning the ligated 18-mer and 30-mer were cut out. RNA was eluted from gel pieces in 800 µl gel elution buffer (0.3 M NaCl, 0.1 % SDS) rotating overnight. To concentrate samples, RNA was precipitated by addition of 2.5-3x volumes of ice-cold EtOH for 1 hour at -20°C, after centrifugation for 30 min at 4°C, RNA pellet was washed using 80% ice-cold EtOH and eluted in 6 µl H2O. **5’ adapter ligation and purification.** 1 µl of 5’ adapter (10 µM) was added and incubated for 5 min at 65°C before addition of T4 ligase buffer (NEB), final 25% PEG8000 and T4 RNA ligase 1 (NEB, M0204L). The ligation reaction was incubated overnight at 4°C. RNA Clean & Concentrator kit (Zymo, R1013) was used for purification and samples were eluted in 12 µl H2O. The structure of the final ligation product is indicated in Fig. S3a. **Reverse transcription and library amplification.** To generate cDNA, 200U Superscript II or III reverse transcriptase (Invitrogen, 18064014/18080044) was used with 40U of RNaseOUT (Invitrogen, 10777019) in a 20 μL volume without heat inactivation. The resulting cDNA was cleaned up using ExoSAP-IT (Applied Biosystems, 78201) enzymatic cleanup. After heat inactivation (15 min at 85°C), the cDNA was used for real-time quantitative PCR (qPCR) with the Kapa HiFi HotStart Library Amp kit (Roche, KK2612). For Illumina short-read sequencing, libraries were amplified using dual indexed TruSeq i5/i7 primers until the fluorescence level reached standard 3 provided with the kit. For unoxidized samples with an initial RNA input of 1.5 μg per library, 11 cycles were commonly used. Amplified libraries were purified using the Zymoclean Gel DNA recovery kit (Zymo, D4008) from a 3% low range agarose gel (BioRad, 1613107) to remove adapter dimers.

### Computational/statistical analysis/data processing

Ready prepared libraries from RKO cells were sequenced using the Illumina HiSeqV4 SR50 mode. Libraries prepared from mouse samples were sequenced on an Illumina NovaSeq SP in SR100 XP mode and on an Illumina NovaSeq S4 in 150 paired-end mode (B cell and PC experiments). In the latter case, only mate1 was further analysed. Sequencing quality control was conducted with fastqc v 0.11.8. Small RNA sequencing reads obtained from RKO time-course samples were mapped to human genome GRCh38 and the microRNA and spike-ins counts were quantified with the annotation from miRbase^8^, using the same NextFlow pipeline as in Dexheimer et al., 2020^9^. Spike-in normalization was performed for miRNA expression levels with unit of amol/µg of total RNA.

For mouse samples, an updated version of this pipeline was used, in more detail: Raw reads were demultiplexed and analysed with a NextFlow pipeline that orchestrated public bioinformatics tools and custom python scripts based on biopython, pysam, pandas, and numpy libraries^10^. Raw reads were parsed and filtered as follows (Fig. S3b-c):

First, we aligned the expected adapter sequence (‘AGATCGGAAGAGCACACGTCT’) to the read sequence using a global sequence alignment with matches scoring +1 and gap opening/extending penalties -2 and - 1 respectively. Start/end gaps were not penalized. The cumulative alignment score was length-normalized, and reads were filtered (‘no_adapter’ reads) by comparing to a minimum score (0.9 in our experiments). Sequences upstream of the aligned adapter were extracted and interpreted as sRBC barcode (5nt) and UMI (6nt). Sequences upstream of the UMI were then extracted and interpreted as small RNA reads. Small RNA reads shorter than 18nt (‘too_short’) and reads where the extracted sRBC barcode did not match the expectation (‘wrong_srbc’, e.g., due-to cross-sample contamination) were filtered. The resulting pre-filtered FASTQ files were then further processed by fastp (v0.20.1) which was configured to first trim 4nt off the read start and then filter reads based on quality (default settings) and a minimum remaining read length of 18nt (see Fig. S4 for example statistics).

We then filtered and counted reads stemming from our 8 different spike-in RNAs (see below) by aligning the spike-in sequences to the read, allowing for one mismatch. On average, we assigned ∼2-3% of reads to the different spike-ins and filtered them from our datasets. Remaining small RNA reads were then mapped to a transcriptome created from MirGeneDB v2.0 annotations using Tailor v1.1^11^. Transcriptomes (FASTA and GFF3 annotation files) were created by extracting contigs from the human (GRCh38) and mouse (mm10) reference genomes using the respective pre-miRNA annotations, where overlapping annotations resulted in single contigs containing all respective annotations. Mappability tracks for visual inspection and QC were calculated with genmap v1.3.0^12^ (parameters k=18, e=2). The minimum prefix matching parameter of Tailor (-l) was set to 18nt. We then created down-sampled BAM files from the full alignments for visual inspection in a genome browser, debugging and QC purposes.

Finally, small RNA reads were counted by a custom python script. For pre-miRNA annotations, we counted strand-specific overlapping reads. For mature miRNAs, we counted only reads for which the 5’end is within +/- 5bp of the annotation 5’end and for which the 3’end does not exceed more that 5bp beyond the annotation. Soft-clipped bases at the 3’ends of the reads were interpreted as tails, extracted, and counted. Counts of multimapping reads were normalized by the number of optimal alignment positions (1/NH). Resulting count tables for mouse samples were further analysed in R using Rstudio v2022.12.0.

We estimated sequencing library sizes from concentration-normalized methylated spike-in read counts and then normalized read counts by these values. For downstream analysis only miRNAs with reads >0.01 (human samples) or >0.5 (mouse samples) in unoxidized samples were considered. For results tables as used for generating graphs shown in this study, see Extended Data Table 1-2. Plots were generated using ggplot2 (v3.3.6) and GraphPad Prism (v10.0.2). Figures were prepared with Adobe Illustrator (v25.4.1).

## Extended Data Tables

Extended Data Table 1 contains all normalized sequencing data. Extended Data Table 2 contains PC miRNAs and miRNAs identified with high confidence in 1%, 0.1% and 0.01% PC samples. Extended Data Table 3 contains all plasmids used in this study as well as DNA/RNA oligo sequences.

## Supplementary Figures

File contains Supplementary Figures 1-5. S1-S2 show uncropped scans of Western Blot images and Northern blot hybridizations. S3-S5 include information about processing of small RNA libraries as described in the Methods part of this study.

## Data availability

Raw sequencing data are available as Zenodo repository (**10.5281/zenodo.10014351**).

## Code availability

Pipelines used for processing of small RNA sequencing data can be found on GitHub under https://github.com/lengfei5/smallRNA_nf/tree/master/dev_sRBC (used for RKO time-course) and under https://github.com/popitsch/pysrna (used for mouse B cell and plasma cell libraries, supports as well samples from human cells).

## Supplementary information

**Supplementary Fig. S1.**
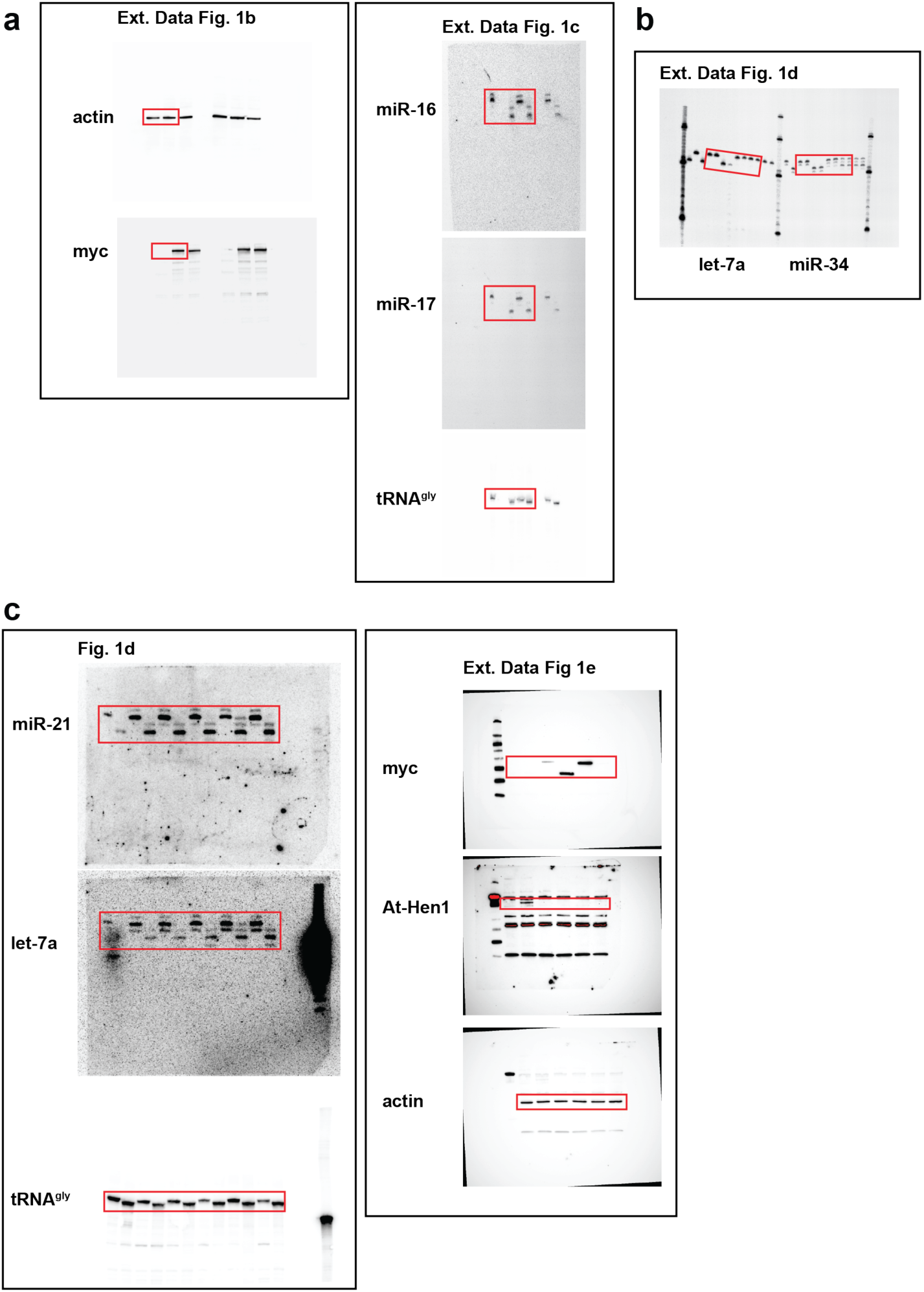
Full scans of Western blots and corresponding Northern hybridizations shown performed with samples obtained from HEK293T cells. **(a)** On the left, WB associated with Extended Data Figure 1b. On the right, Northern hybridization associated with Extended Data Figure 1c. **(b)** In vitro methylation assay associated with Extended Data Figure 1d. **(c)** On the left, Northern hybridization associated with Figure 1d. On the right, WB associated with Extended Data Figure 1e.

**Supplementary Fig. S2.**
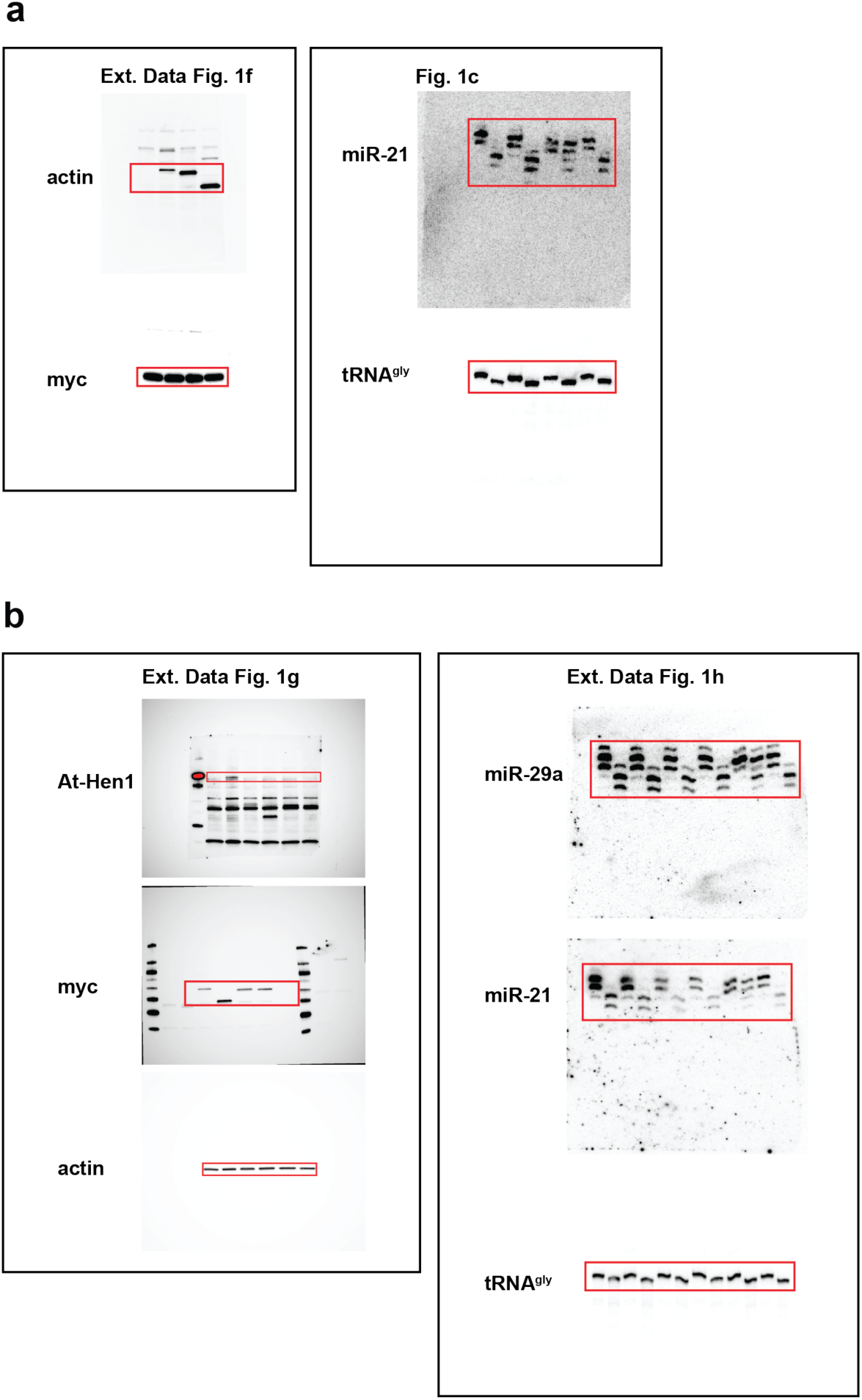
Full scans of Western blots and corresponding Northern hybridizations shown performed with samples obtained from (a) MEF or (b) RKO cells. **(a)** On the left, WB associated with Extended Data Figure 1f. On the right, Northern hybridization associated with Figure 1c. **(b)** On the left, WB associated with Figure 1g. On the right, Northern hybridization associated with Extended Data Figure 1h.

**Supplementary Fig. S3.**
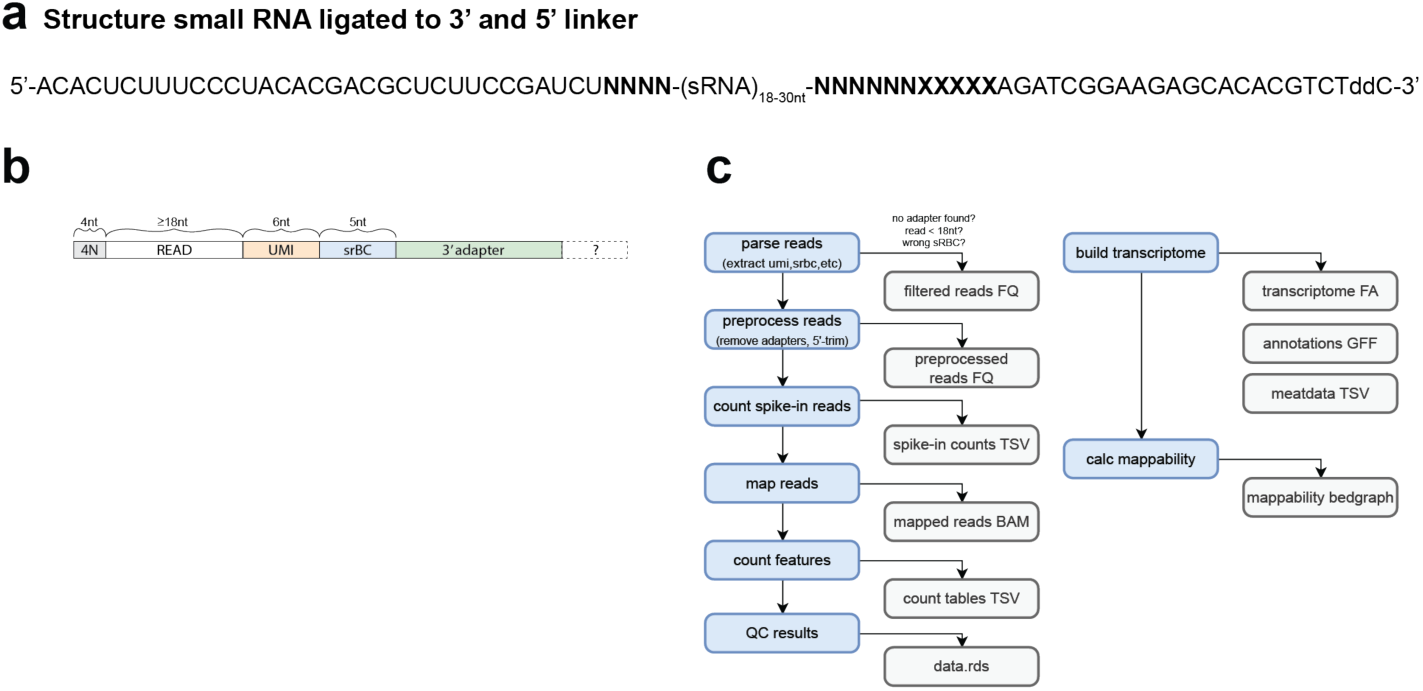
Small RNA libraries additional details for processing. **(a)** Structure of small RNA ligated to 3’ and 5’ linkers. Ns indicate random nucleotides as part of the linkers to counteract ligation bias. XXXXX indicates position of 3’ srBC barcode. **(b)** Outline of sequencing read structure. **(c)** Block diagram providing brief overview of analysis pipeline stages.

**Supplementary Fig. S4.**
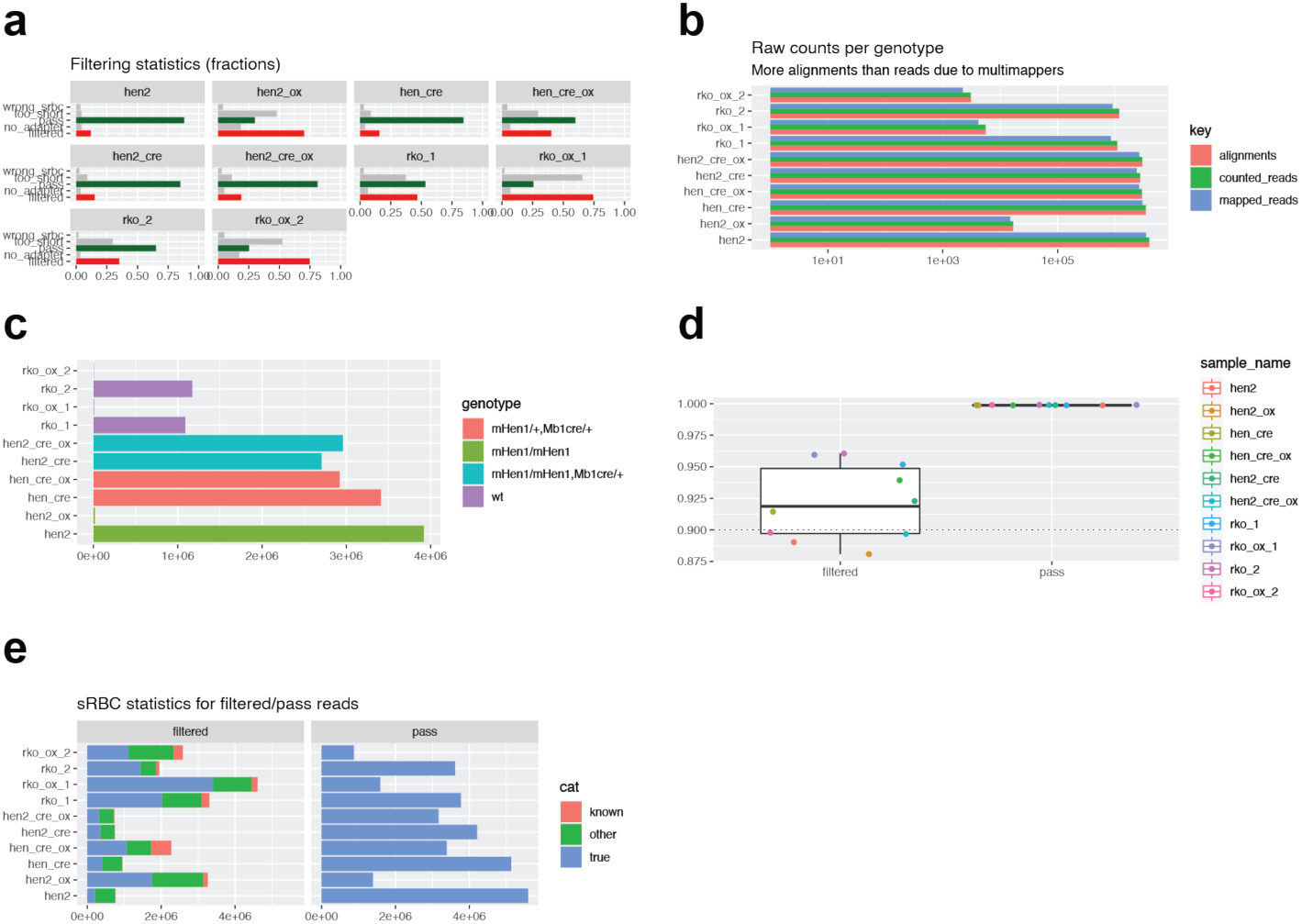
Small RNA pipeline - filter statistics example B cell libraries. **(a-e)** For each genotype oxidized and unoxidized sample is shown. **(a)** Fraction of reads per sample that passed (green) or failed (red) the filtering criteria. Gray bars indicate fractions of filter reasons. **(b)** Number of mapped/counted reads/alignments per sample. X-axis is logarithmic. **(c)** Mapped read counts per sample, colored by genotype. **(d)** Mean adapter alignment scores for reads that passed/failed the filtering criteria; dashed line indicates the configured minimum alignment score threshold. **(e)** Numbers of found srBC sequences for filtered/passed reads. ‘True’: found sRBC matches expected one; ‘Known’: found srBC matches one expected for another sample; ‘other’: found sRBC matches unknown sequence. High numbers of ‘known’ sRBC sequences could reveal sample mix-ups or cross-sample contamination.

